# Innate immune responses yield tissue-specific bottlenecks that scale with pathogen dose

**DOI:** 10.1101/2023.06.09.543079

**Authors:** Karthik Hullahalli, Katherine G. Dailey, Matthew K. Waldor

**Affiliations:** Department of Microbiology, Harvard Medical School, Boston, MA 02115; Division of Infectious Diseases, Brigham & Women’s Hospital, Boston, MA 02115

## Abstract

To cause infection, pathogens must overcome bottlenecks imposed by the host immune system. These bottlenecks restrict the inoculum and largely determine whether pathogen exposure results in disease. Infection bottlenecks therefore quantify the effectiveness of immune barriers. Here, using a model of *Escherichia coli* systemic infection, we identify bottlenecks that tighten or widen with higher inoculum sizes, revealing that the efficacy of innate immune responses can increase or decrease with pathogen dose. We term this concept “dose scaling”. During *E. coli* systemic infection, dose scaling is tissue specific, dependent on the LPS receptor TLR4, and can be recapitulated by mimicking high doses with killed bacteria. Scaling is therefore due to sensing of pathogen molecules rather than interactions between the host and live bacteria. We propose that dose scaling quantitatively links innate immunity with infection bottlenecks and is a valuable framework for understanding how the inoculum size governs the outcome of pathogen exposure.

## Introduction

The COVID-19 pandemic has catalyzed renewed interest in the long-standing question of how the size of a pathogen inoculum impacts subsequent infection (1, 2). A critical parameter that determines whether a given pathogen dose can establish infection is the bottleneck, which represents host processes that eliminate inoculated microorganisms (3–7). Organisms that survive bottlenecks and give rise to the population at a site of infection are known as the founding population (FP)(8). FP cannot be measured by enumerating colony forming units (CFU) alone since total burden is governed by both the infection bottleneck and pathogen replication.

We developed STAMPR, a methodology that quantifies FP with barcoded bacteria (9). The relationship between FP and dose measures the bottleneck (4, 6). The few studies that have measured infection bottlenecks with barcoded bacteria have focused on enteric infections and found that bottlenecks restrict fixed fractions of the inoculum rather than fixed numbers (4, 6). Thus, barriers to enteric colonization, such as stomach acid and the microbiota, have the same fractional efficacy at low or high doses. However, in other infection models, it is possible that bottlenecks constrict or widen in response to changes in dose. For example, positive feedback in the innate immune system may tighten bottlenecks at higher doses, or bottlenecks may widen at higher doses if immune effectors are overwhelmed.

We recently profiled the dynamics of systemic *E. coli* infection in mice (10). Following intravenous (IV) inoculation, multiple interconnected components of the innate immune system, including the production of proinflammatory cytokines and infiltration of immune cells to tissues, are rapidly triggered in a manner dependent on the LPS receptor TLR4. These factors impose a bottleneck and govern infection establishment; FP explicitly quantifies the collective efficacy of these innate immune constituents at a single dose. The dose-FP relationship (i.e., the bottleneck) thus quantifies innate immune potency across all doses. In this study, we leveraged STAMPR to define the broader role of TLR4 in modulating the dynamics of *E. coli* systemic infection and provide a conceptual link between dose, bottlenecks, and innate immunity.

## Results and discussion

### Dose-founding population curves reveal tissue-specific potency of innate immunity

TLR4 knockout mice (TLR4^KO^) mice and heterozygous littermates (TLR4^Het^) were intravenously inoculated with a range of doses of barcoded *E. coli*. Five days post infection (dpi), the lungs, liver, and spleen were harvested for bacterial enumeration and STAMPR analysis. All organs exhibited dose-dependent increases in CFU. At higher doses, abscesses appeared in the livers of TLR4^Het^ animals, but not in TLR4^KO^ animals (Figure 1A, blue box). Consistent with our companion study (11), abscesses result from bacterial replication (low FP and high CFU). TLR4^KO^ mice are resistant to abscess formation (Figure 1A, outside blue box) but have higher FP (Figure 1B), indicating that in the absence of TLR4, more bacteria from the inoculum survive infection bottlenecks. In the spleen, TLR4^KO^ animals also had higher burdens than TLR4^Het^ animals (Figure 1C). FPs were higher in TLR4^KO^ spleens as well, revealing that TLR4 controls splenic infection bottlenecks (Figure 1D). In contrast to both the liver and spleen, neither bacterial burden nor FP was influenced by TLR4 in the lungs (Figure 1E-F).

**Figure 1.**
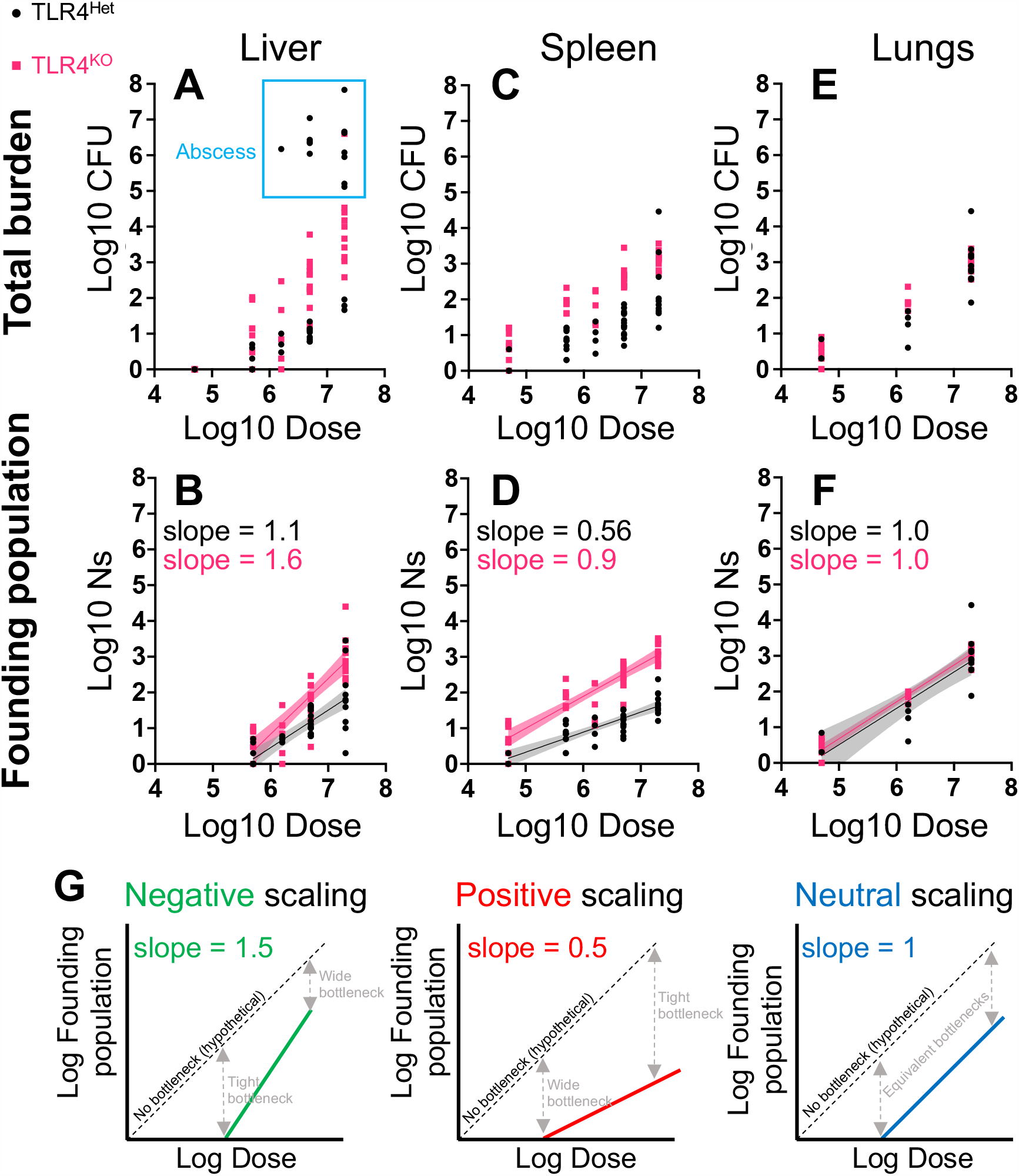
Bottleneck-dose response analysis for *E*. *coli* systemic infection. A-B) CFU (A) and FP (Ns) (B) are shown for the liver as a function of dose. Each point represents an animal and the blue box indicates animals that developed abscesses. C-F are identical to A-B but represent spleen and lung CFU (C and E) and FP (D and F). Best fit lines from linear regression in FP plots are shown with 95% confidence bands. G) Dose-FP plots for different scaling patterns. Dotted lines represent a 0% bottleneck. With slopes greater than one, fractionally more bacteria survive to become founders at higher doses, suggesting that innate immune responses are less effective at higher inoculum sizes, and therefore “negatively scale” with dose. With slopes less than one, the immune response is “positively scaled”, since fractionally fewer bacteria survive (more are killed) at higher doses. With a slope equal to 1, a fixed fraction of the inoculum survives host bottlenecks.

Bottlenecks are quantified by the relationship between FP and inoculum size (4, 6). A slope of 1 on dose-FP curves indicate that numerically more bacteria are eliminated at higher doses, but the fraction killed (and the fraction surviving) is constant. TLR4^KO^ animals had higher dose-FP slopes compared to littermate TLR4^Het^ animals in the spleen (0.9 ± 0.07 (Std. error) vs 0.56 ± 0.06, Figure 1D) and liver (1.6 ± 0.17 vs 1.1 ± 0.14, Figure 1B). Although TLR4 influences the slopes in both organs by a similar magnitude, the specific numerical values reveal tissue-specific differences in the potency of innate immunity. The slope of <1 in the TLR4^Het^ spleen is an example of ‘positive scaling’ (Figure 1G), where innate immunity is more effective at higher inoculum sizes; fractionally fewer bacteria survive to become founders at larger doses (fractionally more are killed). Positive scaling in the spleen is dependent on TLR4, since in its absence, the slope increases to ∼1 (‘neutral scaling’ Figure 1G). Neutral scaling was observed in the livers of TLR4^Het^ animals. In the absence of TLR4 in the liver, the slope increased to >1; fractionally more bacteria survive at higher doses (fractionally fewer are killed). Thus, in the absence of TLR4, the hepatic innate immune response is fractionally less effective at larger inoculum sizes, which we term ‘negative scaling’. In contrast to the liver and spleen, we observed neutral scaling in the lung independent of TLR4. Collectively, these results reveal that in addition to controlling tissue-specific immune responses, TLR4 governs the extent to which the efficacy of these responses quantitatively scale with inoculum size; mice lacking TLR4 have disproportionately reduced potency of innate immune responses at higher doses.

### Dose scaling does not require live bacteria

We hypothesized that dose scaling is separable from the actual number of inoculated organisms and is instead dependent on how the innate immune system senses and ‘interprets’ the inoculum size. To test this idea, we spiked killed bacteria into a live inoculum to simulate some of the immune-stimulatory aspects of high doses. TLR4^Het^ and TLR4^KO^ animals were inoculated either with 1.25×10^6^ CFU live bacteria or the same quantity of live cells plus 5×10^7^ formalin-killed *E. coli* (Figure 2A). We confirmed that both groups received identical amounts of live bacteria (Mean Log10 Dose ± St.dev, control: 6.03 ± 0.14, spike-in: 6.13 ± 0.10). CFU (Figure 2BDF) and FP (Figure 2CEG) were calculated at 5 dpi.

**Figure 2.**
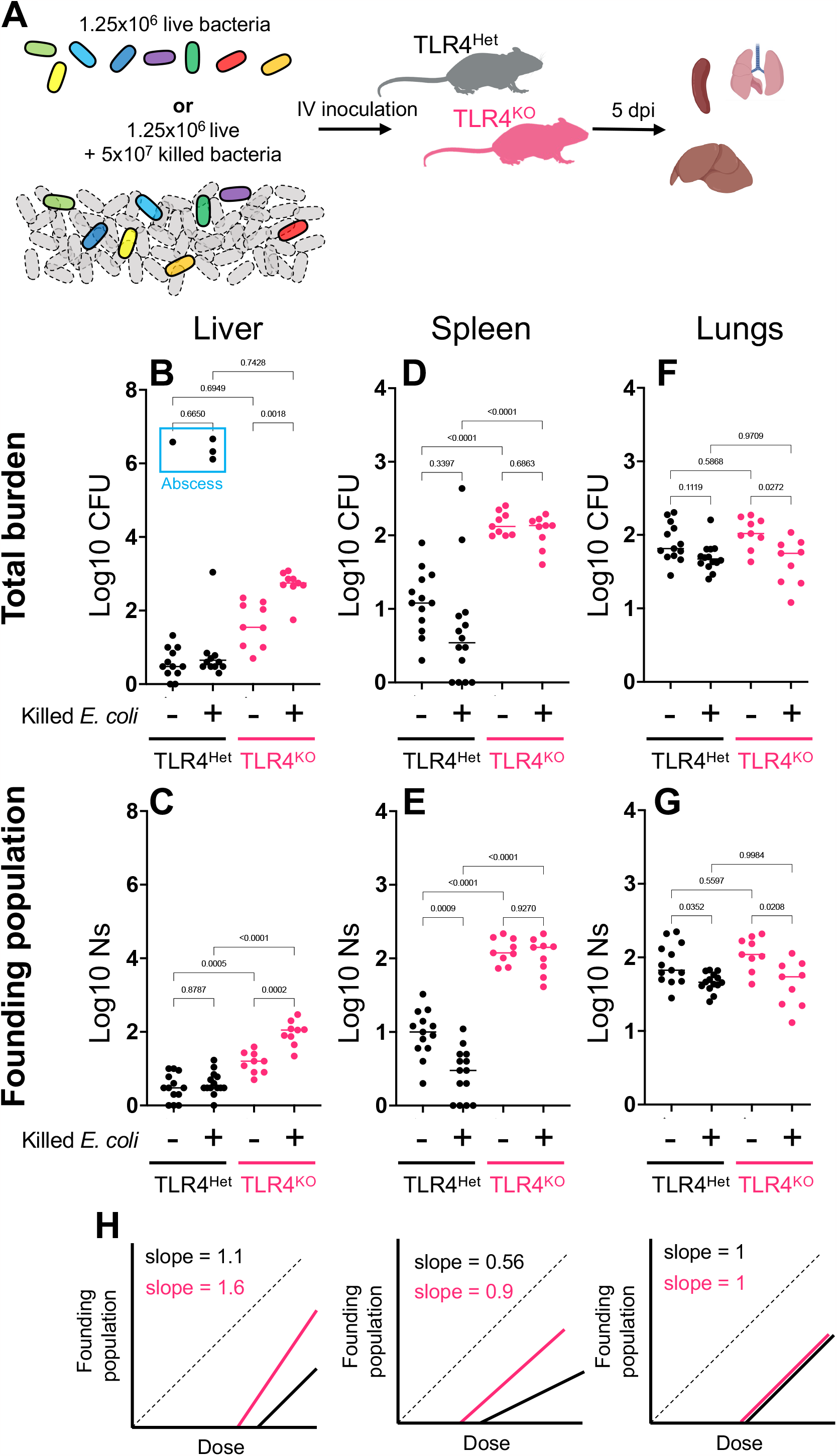
Tissue and TLR4 dependent responses to spike-in of killed bacteria. A) Barcoded live bacteria or the same quantity of live cells plus 40-fold excess of formalin-fixed bacteria were IV inoculated into TLR4^Het^ or TLR4^KO^ littermates. B-C) CFU (B) and FP (Ns) (C) of the liver are shown (lines represent medians). The blue box indicates animals that developed abscesses. D-G are identical to B-C but represent spleen and lung CFU (D and F) and FP (E and G). H) Curves from Figure 1 (left to right: liver, spleen, and lung) are schematized for reference

The potency of the immune response in the spleen was greater at higher doses (positive scaling) in a manner dependent on TLR4 (Figure 1D). Consistent with increased innate immune efficacy at higher doses, spike-in of killed bacteria resulted in 3-fold fewer live founders in TLR4^Het^ spleens (Figure 2E), and the decrease in FP was not observed in the absence of TLR4. In contrast, the livers of TLR4^KO^ animals were scaled negatively (Figure 1B). With spiked killed cells, TLR4^KO^ livers had significantly greater CFU and 5-fold more founders than control animals without spiked-in killed cells (Figure 2C). No changes in CFU or FP were seen with spiked-in cells in the livers of TLR4^Het^ animals, consistent with neutral scaling (Figure 2C). Therefore, positive scaling in the spleen and negative scaling in the liver do not require live bacteria.

The magnitude of changes in FP in response to killed cells is very close to predictions from dose-FP curves (Figure 1). By multiplying the fractional bottleneck (FP/Dose) at a 2 ×10^7^ dose with an actual dose of 1.25×10^6^, the TLR4^Het^ spleen and TLR4^KO^ liver is predicted to have 3-fold fewer and 4-fold greater founders, respectively, relative to control animals without spiked-in cells. Note that the effect of killed cells on CFU in the spleen is not statistically significant despite a significant difference in FP, since two mice had stochastic bacterial replication (Figure 2D). However, the livers of TLR4^KO^ mice had significantly greater CFU after spike-in (Figure 2B). Thus, when bacteria can replicate substantially, FP rather than CFU is a more faithful measure of scaling since it is not confounded by bacterial expansion. Although the lungs exhibited neutral scaling, we observed a minor but statistically significant decrease in CFU and FP when killed cells were spiked in (Figure 2FG).

Together these data show that dose scaling can be recapitulated without increasing the inoculum size. Furthermore, the two seemingly opposite effects of spiking in killed cells (higher FP in TLR4^KO^ livers but lower FP in TLR4^Het^ spleen) result from a similar conceptual basis in dose scaling. The dependence of the effects of killed cell spike-in on TLR4 suggests that dose-scaling results from quantitative changes in the LPS-induced immune response, rather than the actual pathogen dose. The increase in liver FP following spike-in only in the absence of TLR4 may suggest that pathogen molecules other than LPS yield negative scaling patterns. We speculate that the host pathways engaged by these other molecules are more easily exhausted in the absence of TLR4. Other pathogen and host factors that control immune responses, such as TLR5 (flagellin) or Nod2 (peptidoglycan), may benefit from analysis with the dose-FP paradigm to decipher how innate immunity scales with dose to regulate infection outcome.

### Concluding remarks

In this study, we describe the concept of dose scaling, which relates changes in inoculum size with quantitative changes in the efficacy of innate immune responses. We show that different pathogen doses can yield tighter or wider bottlenecks in a tissue-specific manner. Since scaling can in part be recapitulated with killed organisms, leveraging scaling may represent a novel framework to identify therapeutics that tighten infection bottlenecks. We hypothesize that positive and negative scaling arise from differences in the rate of induction of individual components of the innate immune response as a function of inoculum size. Further studies of dose scaling will provide critical contextualization for natural infections where inoculum sizes are not controlled.

## Acknowledgements

This work is supported by NIH F31 AI156949 (K.H.), NIH R01 AI042347 (M.K.W.), and the Howard Hughes Medical Institute (M.K.W.).

## Materials and Methods

### Ethics Statement

All animal experiments were conducted in accordance with the recommendations in the Guide for the Care and Use of Laboratory Animals of the National Institutes of Health and the Animal Welfare Act of the United States Department of Agriculture using protocols reviewed and approved by Brigham and Women’s Hospital Committee on Animals (Institutional Animal Care and Use Committee protocol number 2016N000416 and Animal Welfare Assurance of Compliance number A4752-01)

### Bacterial culture

The *E. coli* strain used in this study was CHS7, derived from strain CFT073 (12). All experiments used a barcoded library of CHS7(10). Bacteria were routinely plated on LB + 50 μg/ml kanamycin agar plates for CFU enumeration.

### Animal Experiments

The TLR4^KO^/TLR4^Het^ colony was established by crossing TLR4^KO^ males (Jackson stock #029015) with C57Bl/6J (Jackson stock #000664) females. TLR4^Het^ female offspring were crossed with TLR4^KO^ males to generate experimental TLR4^KO^ and TLR4^Het^ littermates. Animals were bred in a Biosafety Level 1 helicobacter-free facility at the Brigham and Women’s Hospital. At 7-8 weeks of age, animals were transferred to a Biosafety Level 2 room. At 8-10 weeks old, animals (males and females) were inoculated intravenously with the barcoded library of CHS7. The bacterial inoculum was prepared by thawing small frozen aliquots of a master lot of barcoded CHS7 and diluting directly in PBS. To prepare the inoculum containing killed bacteria, frozen aliquots were diluted in 4% paraformaldehyde and incubated at room temperature for 20 minutes. Cells were centrifuged, washed in PBS, and resuspended in 1ml of PBS, after which live bacteria were added. The 100μl inoculum was delivered intravenously via the lateral tail vein with a 27G needle while animals were restrained in a Broome-style restrainer (Plas Labs). A heating pad was used to facilitate dilation of the tail vein. At 5 days post inoculation, mice were euthanized with isoflurane and cervical dislocation, after which the lungs, spleen, and liver were harvested. Prior to harvesting the liver, the bile was aspirated, and the gallbladder was removed. Organs were homogenized by bead beating with two 3.2mm stainless steel beads for two minutes in PBS.

### STAMPR analysis

1ml of organ homogenates were plated on 150mm agar plates. Note that liver samples were homogenized in a total of 4ml, but only 1ml was plated for STAMPR analysis. To accurately compare CFU and FP values, all liver CFU values reported in this study are for ¼ of the liver. Colonies were picked or lawns were scraped into PBS + 25% glycerol and stored at -80º C.

To amplify the barcode region, cell suspensions were diluted ∼1:50 in water, or up to 1:10 if cell density was low (i.e., few CFU in the organ). Diluted suspensions were boiled (95ºC for 15 min) and 3μl was used for PCR as previously described, with minor modifications (6). PCR reactions were performed with 2x OneTaq HotStart DNA Polymerase master mix (New England Biolabs), using 6 μl of forward and reverse primer (1μM each). The forward primers were designed such that the first nucleotides to be sequenced were a region of variable length to facilitate optimal color balancing. These variability regions also contained sequence diversity for multiplexing and demultiplexing. Reverse primers contained i7 indexes from the TruSeq DNA PCR Free kit (Illumina). PCR reactions were run on agarose gels to verify amplification, pooled, and then purified using the GeneJet PCR purification kit. Amplicons were quantified with Qubit and sequenced as 1 x 78 nt reads on MiSeq or NextSeq 1000 instruments (Illumina)

Reverse primer demultiplexing with i7 was performed on BaseSpace (Illumina), and forward primer demultiplexing was performed using custom R scripts. Demultiplexed reads (.fastq files) were trimmed and mapped to a previously defined list of reference barcodes using CLC genomics workbench. Read count tables, where columns specify the sample and rows specify the barcode, were exported as .csv files and used for subsequent analysis. For samples sequenced on a NextSeq 1000, we included an additional index hopping correction. For each sample *n*_*0*_, we identified every other sample *n*_*i*_ that was separated by only one index. Barcode counts were combined for all *n*_*i*_ samples, multiplicatively scaled, and subtracted from sample *n*_*0*_. No more than 3% of reads were removed from any sample.

Founding population size was approximated by calculation of Ns, which is defined as the number of times the input library must be sampled from a multinomial distribution to observe a given number of barcodes. Ns calculation was performed as previously described(6, 13) . Briefly, read count tables were first converted to frequencies. To identify noise thresholds, we first identified large gaps in barcode frequency between sorted barcodes (greater than 10-fold) as well as breaks that delineate sub-populations as defined by the STAMPR algorithm (described extensively in (9)). All barcodes below the frequency threshold were defined as noise and set to 0 reads. Ns was calculated by first resampling the input library to the read depth of the output sample. The resulting distribution was then iteratively resampled from 1 to ∼10,000 times. A sampling size of ∼10,000 is sufficient to detect all ∼1000 barcodes in the library and defines the maximum resolution of Ns. A “reference resample curve” was generated, where the x-axis is the number of times sampled (from 1 to ∼10,000) and the y-axis is the number of barcodes detected. The number of barcodes in each sample after noise thresholding was used as input for inverse interpolation from the reference resample curve to calculate Ns.

### Statistics

Linear regression in Figure 1 was performed in GraphPad Prism. This study assumes that curves are linear, but it is possible that with a substantially larger number of replicates a nonlinear fit would be a more appropriate approximation of the dose-FP trend (4). The 5 x 10^4^ dose was excluded from linear regression analysis in the liver, since 7 animals had 0 CFU and 4 animals had 1 CFU. Statistical significance in Figure 2 was assessed using Brown-Forsythe and Welch ANOVA tests followed by Dunnett’s T3 multiple comparisons test (GraphPad Prism).

